# Two novel strains of *Torulaspora delbrueckii* isolated from the honey bee microbiome and their use in honey fermentation

**DOI:** 10.1101/264317

**Authors:** Joseph P. Barry, Mindy S. Metz, Justin Hughey, Adam Quirk, Matthew L. Bochman

## Abstract

Yeasts are ubiquitous microbes found in virtually all environments. Many yeast species can ferment sugar into ethanol and CO_2_, and humans have taken advantage of these characteristics to produce fermented beverages for thousands of years. As a naturally abundant source of fermentable sugar, honey has had a central role in such fermentations since Neolithic times. However, as beverage fermentation has become industrialized, the processes have been streamlined, including the narrow and almost exclusive usage of yeasts in the genus *Saccharomyces* for fermentation. We set out to identify wild honey- or honey bee-related yeasts that can be used in honey fermentation. Here, we isolated two strains of *Torulaspora delbrueckii* from the gut of a locally collected honey bee. Both strains were able to ferment honey sugar into mead but failed to metabolize more than a modest amount of wort sugar in trial beer fermentations. Further, the meads fermented by the *T. delbrueckii* strains displayed better sensory characteristics than mead fermented by a champagne yeast. The combination of *T. delbrueckii* and champagne yeast strains was also able to rapidly ferment honey at an industrial scale. Thus, wild yeasts represent a largely untapped reservoir for the introduction of desirable sensory characteristics in fermented beverages such as mead.

## Introduction

Mankind has been fermenting foods and beverages for thousands of years [1–4], which certainly predates our knowledge that yeasts were the microbial agents responsible for metabolizing sugar into alcohol. Honey, which is produced by honey bees such as *Apis mellifera*, is a natural source of abundant, readily fermentable sugar and has been found as an ingredient in some of the earliest known fermented beverages [2]. When a dilute honey solution is fermented on its own without the addition of fruit/fruit juice or other additives, this creates a traditionally strong (8-18% alcohol by volume, ABV) beverage called mead (reviewed in [5]).

Much like craft brewing, mead making is currently experiencing a renewed interest at the amateur and professional levels worldwide. Indeed, one average, a new meadery opens in the United States every 3 days, and one opens every 7 days in the rest of the world [6]. With this proliferation of mead making has come experimentation with the traditional methods, including aging mead in oak or previously used spirit barrels and fermentation with non-*Saccharomyces* microbes to generate sour and “funky” meads [6]. In many ways, this again mirrors the craft brewing scene and current popularity of sour beer [7–9].

Another factor driving innovation in fermented beverage production is the “local movement” [10, 11], i.e., the push to use locally sourced ingredients. In the case of mead, it is often relatively simple to utilize locally produced honey, but few meaderies use indigenous microbes unless they rely on those naturally found in the honey itself. Such microbes are naturally resistant to high osmotic stress from concentrated sugar in honey and typically come from pollen/flower, bee gut microbiome, and dirt/dust contamination of the honey (reviewed in [12, 13]). These organisms include multiple species from dozens of different genera of bacteria (e.g., *Bacillus, Citrobacter*, and *LactoBacillus*), yeast (e.g., *Saccharomyces, SchizoSaccharomyces*, and *Pichia*), and molds.

Here, we describe the bio-prospecting for novel yeasts to use in honey fermentation. We concentrated on ultra-local strains by attempting to enrich for ethanol-tolerant yeasts in honey, the hive, and honey bees themselves. Ultimately, we recovered two strains of the yeast *Torulaspora delbrueckii* from the honey bee microbiome and characterized them for their ability to ferment honey. Many *T. delbrueckii* strains have previously been examined for their fermentative abilities and were found to improve wine quality (see [14] and references therein) and create distinct beer flavors in beer [15–17]. Compared to *S. cerevisiae, T. delbrueckii* is generally found to generate lower levels of ethanol [16] but a wider range of fruity aromas [18, 19]. However, in many of these cases, *T. delbreuckii* is used in combination with *S. cerevisiae* for sequential fermentations [14] or mixed culture fermentations [15].

Currently, there are no reports of *T. delbrueckii* being used to make mead and how this species may impact the sensory qualities of the final product. Here, we found that compared to a traditional *Saccharomyces* cerevisiae control strain (WLP715), the mead produced by the *T. delbrueckii* strains YH178 and YH179 scored more favorably during sensory analysis. Further, *T. delbrueckii* YH178 and YH179 were successfully used in mixed honey fermentations with *S. cerevisiae* WLP715 at a production scale. These data suggest that *T. delbrueckii* YH178 and YH179 have beneficial honey fermentation profiles and are suitable for mead production by homebrewers and professional mead makers.

## Results and Discussion

### Isolation of T. delbrueckii strains YH178 and YH179

We initially set out to isolate ethanol-tolerant yeasts from multiple bee-related sources. Samples of honey, honeycomb, propolis, and fragments of a wooden beehive were surveyed for yeasts, but no microbes were recovered using our wild yeast enrichment protocol [7]. This failure was likely not due to the stringency (*e.g*., high ethanol and osmotic stress tolerance) of the method used because we previously recovered hundreds of wild ethanol-tolerant yeasts from a variety of environmental sources. However, as only small volumes of samples were surveyed from a single hive, a more extensive search may have yielded suitable yeasts. Alternatively, the lack of ethanol-tolerant yeasts may be attributed to the well-characterized hygienic behavior of honey bees, which are known to keep their hives clean in order to avoid disease [20–23].

We next focused our bio-prospecting efforts on the bees themselves. We again failed to isolate ethanol-tolerant yeast from the outer surface of several bees *via* a quick rinse with yeast enrichment medium or prolonged incubation of a bee submerged in the same medium at 30°C with aeration. However, when a bee was submerged in yeast enrichment medium and its abdomen was ruptured, microbial growth was visible in the culture after 24 h. Plating for single colonies on WLN agar revealed that two distinct strains of yeast were present, which differed in color (Fig. 1A). These strains were designated YH178 (darker green on WLN agar) and YH179 (pale green) as part of our larger yeast hunting efforts and were identified as *T. delbrueckii* isolates by rDNA sequencing (Fig. 1B). For comparison, the *S. cerevisiae* strain WLP715 is shown, which attains an even lighter shade of green than YH179 on WLN agar (Fig. 1A)

**Figure 1.**
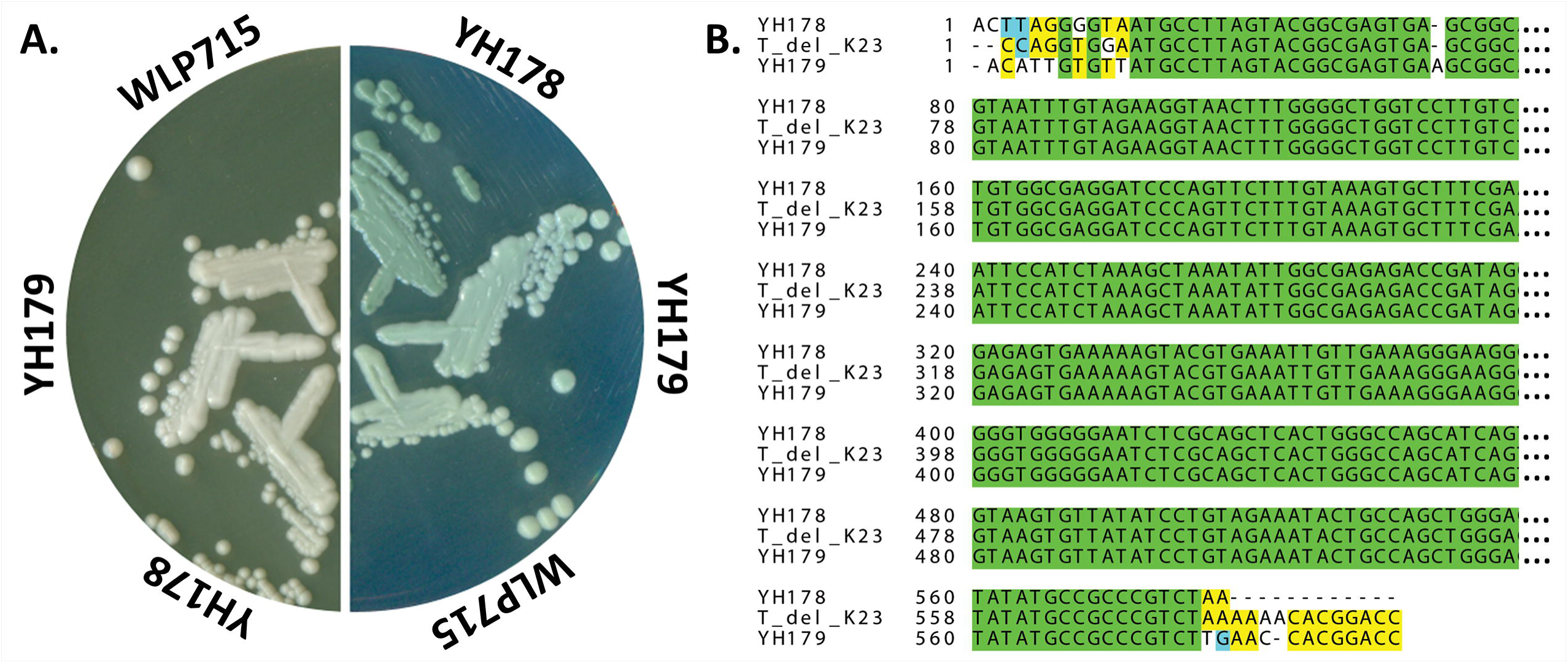
YH178 and YH179 are both strains of *T. delbrueckii*. A) Colony morphology of YH178, YH179, and *S. cerevisiae* WLP715 on YPD (left) and WLN (right) agar plates. The indicated strains were streaked for single colonies and grown at 30°C for 2 days prior to capturing images of the plates with a flatbed scanner. B) Sequence alignments revealed that YH178 and YH179 are isolates of *T. delbrueckii*. A portion of the 26S rDNA of YH178 and YH179 was PCR-amplified and sequenced, and the sequences are aligned to the closest related strain in the NCBI nucleotide database. T_del_K23 denotes the 26S rDNA gene of *T. delbrueckii* strain K23 (GenBank accession KP852445.1). The ellipsis at the end of each line represents the missing sequence that was truncated due to spatial constraints. The full alignment is shown in Supplemental Figure S1. Green, fully conserved; yellow, conserved among two sequences; blue, similarity (pyrimidines); and dashes, sequence gaps due to indels.

Although no colony size variation was noted on either YPD or WLN plates at the end of a 2-day incubation (Fig. 1A), YH178 attained this colony size faster than YH179 when observed prior to the end point of growth (data not shown). This suggested that YH178 has a faster doubling time than YH179. However, plotting growth curves for these strains grown in YPD medium revealed that YH178 had a shorter lag phase (< 4 h; Fig. 2A) to exponential growth than YH179 (> 6 h; Fig 2B). Otherwise, these strains displayed similar doubling times and final cell densities.

**Figure 2.**
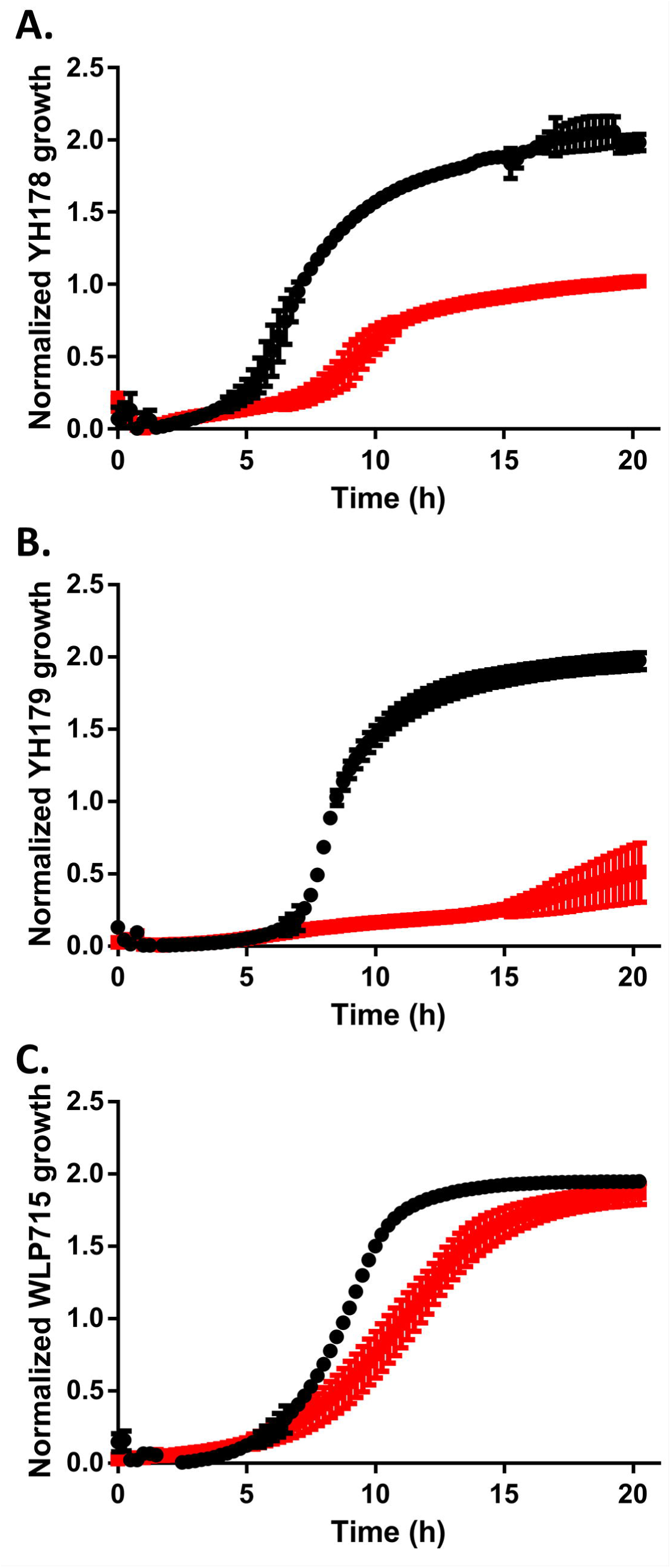
YH178 and YH179 grow poorly in the presence of maltose. YH178 (A), YH179 (B), and *S. cerevisiae* WLP715 (C) were grown to saturation overnight in YPD medium, back-diluted into YPD (back) or YPM (red) medium, and grown overnight in a 96-well plate. Growth was monitored by recording the OD_660_ of each culture and is plotted vs. time. The plotted points are the averages of three independent experiments, and the error bars are the standard deviations.

### YH178 and YH179 can use honey as a carbon source during fermentation

To determine if YH178 and/or YH179 could ferment honey, we inoculated a diluted honey solution with an equivalent number of cells from the *T. delbrueckii* strains singly and in combination. As a positive control for fermentation, we also used *S. cerevisiae* WLP715, which is marketed as an appropriate yeast for wine, mead, and cider fermentations [24]. The fermentations were monitored for 20 days, and the results are shown in Figure 3A. At approximately 12 h after inoculation, all cultures displayed signs of fermentation, and WLP715 quickly attenuated the honey solution to a final alcohol by volume (ABV) of 11.5% in 2 weeks (Table 1). The *t*_1/2_ (time necessary for 50% attenuation) was < 2 days.

**Table 1.** Experimental fermentations with *Saccharomyces* cerevisiae, *Torulaspora delbrueckii*, and *Hanseniaspora vineae* strains.

|  | Honey fermentation<sup>1</sup> |  |  | Beer fermentation |  |  |
| --- | --- | --- | --- | --- | --- | --- |
| Strain | Final gravity | ABV<sup>2</sup> | Final pH<sup>3</sup> | Final gravity | ABV | Final pH |
| *S. cerevisiae* WLP001 | N.D. | N.D. | N.D. | 1.011 | 4.33% | 4.40 |
| *S. cerevisiae* WLP715 | 0.992 | 11.5% | 4.25 | N.D. | N.D. | N.D. |
| *T. delbrueckii* YH178 | 0.992 | 11.5% | 4.15 | 1.041 | 0.39% | 5.40 |
| *T. delbrueckii* YH179 | 0.996 | 11.0% | 4.23 | 1.042 | 0.26% | 5.40 |
| YH178 + YH179 | 0.994 | 11.2% | 4.20 | N.D. | N.D. | N.D. |
| *H. vineae* YH72 | 1.018 | 8.2% | 3.31 | 1.016 | 3.68% | 3.26 |

<sup>1</sup> The original gravity of the honey solution was 1.080, and that of the wort was 1.044.
<sup>2</sup> Abbreviations: ABV, alcohol by volume; and N.D., not determined.
<sup>3</sup> The initial pH of the honey solution was 4.62, and that of the wort was 5.43.

**Figure 3.**
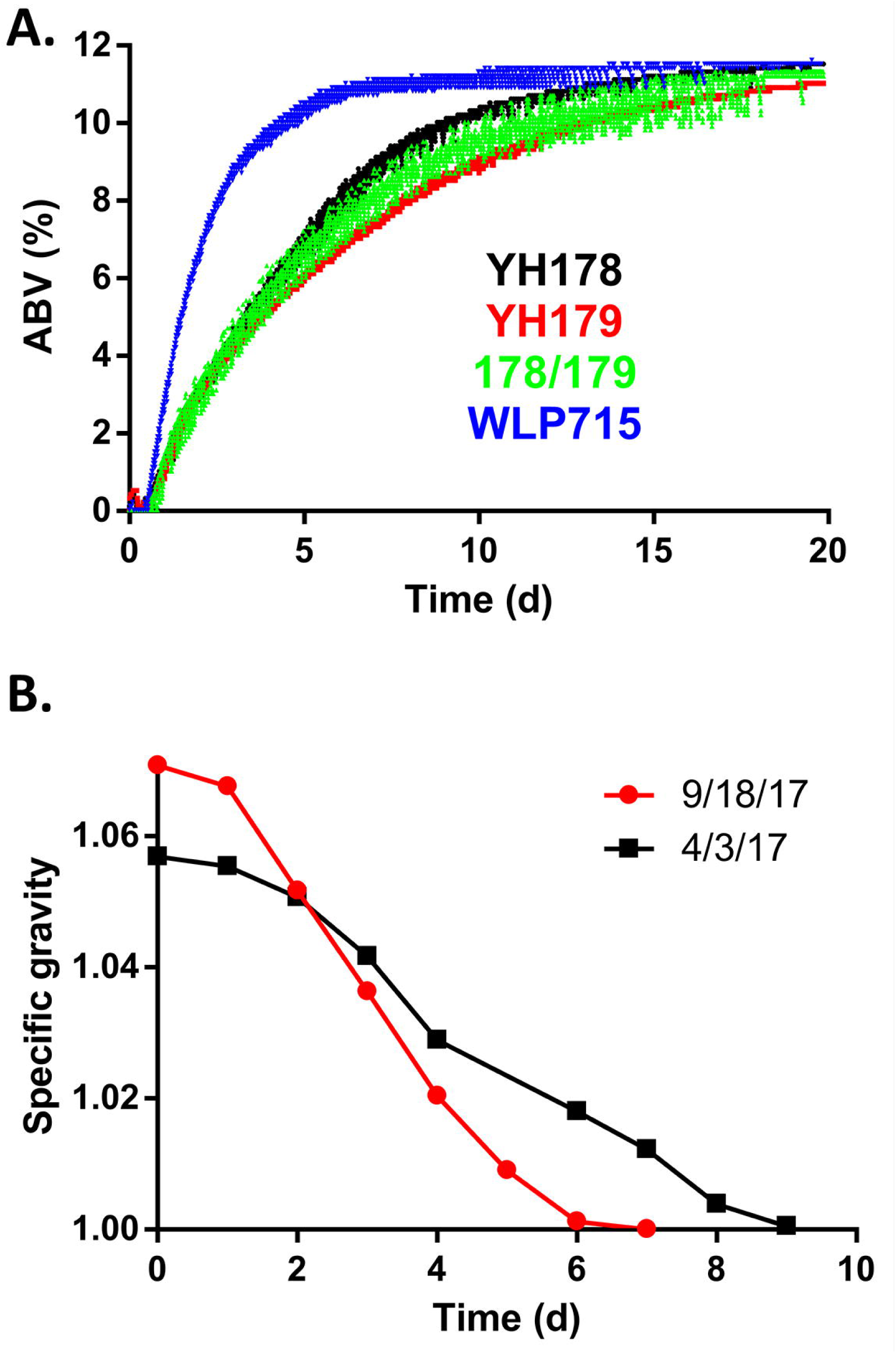
The *T. delbrueckii* isolates can ferment honey into mead. A) The indicated strains were used to inoculate a honey solution, and fermentation of the honey was monitored with a digital hydrometer. The amount of alcohol produced is plotted vs. time. These results are representative of three independent fermentations for each strain or combination of strains (178/179 = a 1:1 mixture of YH178 and YH179 cultures). B) Production-scale honey fermentations using a combination of YH178, YH179, and WLP715 yeasts. These fermentations were conducted in April (4/3/17) and September (9/18/17) of 2017 under similar conditions but with slightly different starting gravities of the honey solution.

In contrast, YH178, YH179, and the combination thereof fermented the honey solution at a much more modest pace. The *t*_1/2_ for these fermentations was ∼4-4.5 days, and while the final ABV for YH178 was identical to that of WLP715 (11.5%), the YH179 and YH178/YH179 combination fermentations only reached 11 and 11.2%, respectively (Table 1). Subsequent experiments did, however, reveal that additional incubation time of the YH179 and YH178/YH179 cultures led to the same terminal ABV of 11.5% by 35 and 28 days, respectively (data not shown).

It is perhaps unsurprising that gut microbiota from the honey bee can metabolize honey, as this is a food source for *A. mellifera* itself. Honey is mainly composed of the hexose monosaccharides glucose and fructose, and most yeast species can utilize one or both as carbon sources for cellular metabolism [25]. Conversely, unadulterated honey contains no maltose [26], which is a glucose disaccharide and the primary sugar in beer wort [27]. We tested YH178 and YH179 for their ability to ferment wort and found that attenuation of the beer was extremely poor after 2 weeks of fermentation (Table 1). In contrast, the *S. cerevisiae* WLP001 ale yeast attenuated to expected levels. The basal level of fermentation displayed by YH178 and YH179 is likely attributable to the *T. delbrueckii* cells consuming the limited quantities of glucose, fructose, and/or sucrose found in typical worts [27]. Indeed, growth curves in YP medium supplemented with 2% maltose (YPM medium) rather than glucose (YPD medium) demonstrate that YH178 grows much more poorly in the YPM than YPD medium (Fig. 2A). Similarly, YH179 displayed only minimal growth in YPM medium (Fig. 2B). The WLP715 strain is able to utilize maltose though, showing only a modest decrease in growth rate in YPM medium compared to YPD medium and no reduction in final cell density (Fig. 2C). While others have successfully isolated yeasts from wasps [28] that can ferment wort into beer, not all yeasts from hymenoptera gut microbiomes are capable of this feat. Thus, individuals interested in isolating yeasts for a particular fermentation milieu are cautioned to select for crucial phenotypes (*e.g*., maltose utilization) early in the bio-prospecting process.

### YH178 and YH179 produced superior meads compared to WLP715

We next compared the sensory characteristics of the meads produced via the above fermentations. This analysis was performed by 10 panelists using a method developed to quantify eight organoleptic qualities of mead [29] on a 20-point scale. Table 2 shows the results of our sensory analysis of mead fermented with YH178, YH179, the combination of YH178 and YH179, and *S. cerevisiae* WLP715. Overall, the meads fermented with YH178 or YH179 were scored more highly than that produced by WLP715 (though not significantly preferred, P ≥ 0.052). The WLP715 mead was notable for its very forward ethanol heat, which was more pleasantly masked by YH178 and YH179 despite having comparable ABVs. However, the sensory panel also detected obvious differences in the meads made with YH178 and YH179. For instance, the aroma of the YH178 mead was more reminiscent of honey than YH179 mead, which some panelists reported as “funky” or simply lacking in honey character. This again corroborates our findings that YH178 and YH179 are distinct strains of *T. delbrueckii*. The mead fermented with the combination of YH178 and YH179 scored highly on aroma but had a grassy flavor that some panelists did not enjoy. Regardless, all of these meads received a “standard” score (12-16 points) [29], indicating that YH178 and YH179 are suitable for home and commercial brewing of mead.

**Table 2.** Sensory evaluation of mead samples.

| Yeast<sup>1</sup> | Appearance | Aroma | Acidity | Balance | Body | Flavor | Finish | Overall quality | Total |
| --- | --- | --- | --- | --- | --- | --- | --- | --- | --- |
| YH178 | 2 ± 0<sup>2</sup> | 3.7 ± 0.9 | 1 ± 0.5 | 1 ± 0.5 | 0.9 ± 0.7 | 1.5 ± 0.3 | 1.3 ± 0.5 | 1.4 ± 0.2 | 12.8 ± 2.4 |
| YH179 | 2 ± 0 | 3.7 ± 1.2 | 1.1 ± 0.7 | 1.7 ± 0.4 | 1.4 ± 0.5 | 1.8 ± 0.4 | 1.8 ± 0.4 | 1.8 ± 0.4 | 15.3 ± 2.4 |
| 178/179 | 2 ± 0 | 4.5 ± 1.2 | 0.8 ± 0.6 | 0.9 ± 0.2 | 0.8 ± 0.6 | 1 ± 0.5 | 1.1 ± 0.2 | 1.2 ± 0.2 | 12.3 ± 1.9 |
| WLP715 | 2 ± 0 | 4.3 ± 0.8 | 0.8 ± 0.6 | 1.3 ± 0.4 | 0.6 ± 0.2 | 1.2 ± 0.6 | 1.1 ± 0.4 | 1.4 ± 0.3 | 12.7 ± 2.2 |
| YH72 | 2 ± 0 | 5.2 ± 0.7 | 1.5 ± 0 | 2 ± 0 | 0.7 ± 0.2 | 2 ± 0 | 2 ± 0 | 2 ± 0 | 17.4 ± 0.8 |

<sup>1</sup> YH178 and YH179 are *Torulaspora delbrueckii* isolates, 178/179 is a mixture of both *T. delbrueckii* strains, WLP715 is a *Saccharomyces cerevisiae* strain, and YH72 is a lactic acid-producing isolate of *Hanseniaspora vineae* [8].
<sup>2</sup> The values presented are the averages ± standard deviations.

Knowing that acidity would be ranked, we also fermented mead with *Hanseniaspora vineae* strain YH72 (Table 2), which produces lactic acid during fermentation [8], as an additional control. As in beer fermentation, the final pH of the H. vineae YH72 mead dropped to pH = 3.31 due to the lactate created by this strain. Surprisingly, during our sensory analysis, the YH72 mead was ranked highest in all categories except one (Body) and achieved a quality score defining it as a “superior” (16-20 points) mead [29] (P ≤ 0.017 vs. all other meads). This was in part due to its assertive peach aroma, pleasant acidic finish, and good balance. It should, however, be noted that the final gravity of this mead was only 1.018, yielding an ABV of 8.2%. Thus, comparing it to dryer meads across all sensory categories can be misleading. Indeed, previous large-scale sensory analysis of sweet and dry meads indicates that consumers prefer sweet meads [30].

### YH178 and YH179 were used in the creation of Honey Schnapps

Based on the performance of YH178 and YH179 in mead fermentation, we sought to determine how well these strains functioned in production-scale (> 20 hL) fermentations. Two large, independent fermentations of diluted honey were conducted as described in the Materials and Methods. A mixed yeast culture containing approximately equal cell counts of YH178, YH179, and WLP715 was used in both cases. As shown in Figure 3B, this yeast blend was able to ferment the solution to a specific gravity of ̃1.000 in < 10 days during both trials. This is consistent with the kinetics of fermentation displayed by these strains individually in laboratory-scale fermentations (Fig. 3A), indicating that YH178 and YH179 can be used for industrial honey fermentations without any additional “domestication” or adaptation, at least in the presence of WLP715. Future experiments with YH178 and YH179 alone are planned, as well as trials of serial re-pitching of these strains into fermentations to select for an increased fermentation rate. Indeed, the desirable rapidity with which WLP715 ferments honey is one of the reasons that it was added to the production-scale fermentations performed in this work. Regardless, the fermented honey solutions were distilled into a schnapps that was back-sweetened with honey and were well received by consumers [31].

## Materials and Methods

### Strains and culture conditions

*T. delbrueckii* strains YH178 and YH179 were supplied by Wild Pitch Yeast, LLC (Bloomington, IN), and *S. cerevisiae* strains WLP001 and WLP715 were purchased from White Labs (San Diego, CA). All strains were routinely grown on yeast extract, peptone, and dextrose (YPD; 1% (w/v) yeast extract, 2% (w/v) peptone, and 2% (w/v) glucose) plates containing 2% (w/v) agar at 30°C and in YPD liquid culture at 30°C with aeration unless otherwise noted. Wallerstein Laboratories nutrient (WLN) agar contained 4 g/L yeast extract, 5 g/L tryptone, 50 g/L glucose, 0.55 g/L KH_2_PO_4_, 0.425 g/L KCl, 0.125 g/L CaCl_2_, 0.125 g/L MgSO_4_, 2.5 mg/L FeCl_3_, 2.5 mg/L MnSO_4_, 22 mg/L bromocresol green, and 15 g/L agar. All strains were stored as 15% (v/v) glycerol stocks at −80°C. Media components were from Fisher Scientific (Pittsburgh, PA, USA) and DOT Scientific (Burnton, MI, USA). All other reagents were of the highest grade commercially available.

### Yeast isolation

Yeasts YH178 and YH179 were isolated from a honey bee sample as described [7]. Briefly, honey bees were collected into sterile glass jars and stored at 4°C before being processed in the laboratory. The bee containing YH178 and YH179 was submerged in 5 mL YPD8E5 (1% yeast extract, 2% peptone, 8% glucose, and 5% (v/v) ethanol), crushed with a sterile pipet, and the culture was incubated at 30°C with aeration for 24 h. A small volume of the culture was then streaked onto a WLN agar plate and for 36-48 h at 30°C. To isolate pure strains, single colonies were picked and restreaked onto fresh WLN agar plates until a uniform colony morphology was obtained.

### Strain identification

YH178 and YH179 were identified as *T. delbrueckii* using the procedure described in [8]. Briefly, 100 μL of saturated overnight culture was mixed with an equal volume of lysis solution (0.2 M LiOAc and 1% SDS) and incubated in a 65°C water bath for ≥15 min to lyse the cells. The genomic (gDNA) was precipitated with 300 μL of 100% isopropanol and vortexing, and it was pelleted with cell debris for 5 min at maximum speed in a microcentrifuge. The supernatant was removed, and the gDNA was resuspended in 50 μL TE buffer (10 mM Tris-HCl, pH 8, and 1 mM EDTA). The variable D1/D2 portion of the eukaryotic 26S rDNA was then amplified by PCR from the gDNA templates using oligos NL1 (GCATATCAATAAGCGGAGGAAAAG) and NL4 (GGTCCGTGTTTCAAGACGG) [32] and the following cycling conditions: 98°C for 5 min; 35 cycles of 98°C for 30 s, 55°C for 30 s, and 72°C for 30 s; and 72°C for 10 min. The amplified DNA was purified using a PCR Purification Kit (Thermo Scientific, Waltham, MA) and sequenced by ACGT, Inc. (Wheeling, IL) using primer NL1. The sequence was used to query the National Center for Biotechnology Information (NCBI) nucleotide database with the Basic Local Alignment Search Tool (BLAST; http://blast.ncbi.nlm.nih.gov/Blast.cgi?CMD=Web&PAGE_TYPE=BlastHome) to identify the most closely related species.

### Growth curves

The yeast strains were grown by inoculating 5 mL YPD liquid medium with single colonies from YPD plates and incubation overnight at 30°C with aeration. The optical density at 660 nm (OD_660_) of each culture was determined using a Beckman Coulter DU730 UV/Vis Spectrophotometer. Then, the cells were diluted to an OD_660_ ∼ 0.1 in 200 μL YPD medium or YPM medium (1% (w/v) yeast extract, 2% (w/v) peptone, and 2% (w/v) maltose) in round-bottom 96-well plates, overlaid with 50 μL mineral oil to prevent evaporation, and incubated at 30°C with shaking in a BioTek Synergy H1 plate reader. The OD_660_ of every well was measured and recorded every 15 min for ∼20 h, and the normalized values (i.e., OD_660_ reads minus the initial OD_660_ value) were plotted vs. time to generate growth curves. All growth experiments were repeated three times, and the plotted points represent the average OD_660_ values (error bars represent the standard deviation).

### Laboratory-scale honey fermentation

Wild flower honey from Fort Wayne, IN was provided by Creek Ridge Honey, LLC and Cardinal Spirits. The honey was diluted approximately 1:5 with sterile deionized water and YP medium to a gravity of 1.080 and final concentrations of yeast extract and peptone of 0.25 and 0.5% (w/v), respectively. Fermentations of this honey solution were performed as described in [8]. Briefly, the yeast strains were grown to saturation (∼200 x 10^6^ cells/mL) in 50 mL of YPD liquid medium at 30°C with aeration and used to inoculate ∼800 mL of honey solution (1.080 original gravity) in 1-L glass cylinders (30 cm tall, 7.5 cm inner diameter). Control fermentations were likewise set up in parallel using the same reagents but inoculating with 50 mL (∼200 x 10^6^ cells/mL) WLP715. These fermentations were incubated at an average temperature of 23.4 ± 0.3°C, and the gravity and alcohol by volume (ABV) were monitored in real time using BeerBug digital hydrometers (Sensor Share, Richmond, VA) for 4 weeks.

### Laboratory-scale wort fermentation

Laboratory-scale wort fermentations were performed as described [8]. Briefly, blonde wort was prepared by mashing 65.9% Pilsner (2 Row) Bel and 26.9% white wheat malt at 65°C (149°F) for 75 min in the presence of 1 g/bbl CaCO_3_ and 1.67 g/bbl CaSO_4_. During the subsequent boil, 7.2% glucose and Saaz hops (to 25 international bittering units) were added. The original gravity (OG) of this wort was 1.044. The yeasts were grown to saturation (∼200 x 10^6^ cells/mL) overnight at 30°C with aeration in 5 mL YPD medium, and these starter cultures were used to inoculate approximately 400 mL of blonde ale wort in glass bottles capped with rubber stoppers and standard plastic airlocks. Control fermentations were set up identically but inoculated with 5 mL (∼200 x 10^6^ cells/mL) WLP001 culture. The fermentation cultures were incubated at room temperature for 2 weeks, and their final gravity was measured using a MISCO digital refractometer (Solon, OH).

### Mead sensory analysis

Sensory analysis was performed using the method developed by the University of California, Davis as described by [29]. Briefly, each mead was sampled in a blinded fashion by 10 or more individuals and immediately scored for appearance (2 points), aroma (6 points), and acidity, balance, body, flavor, finish, and overall quality (2 points each). Using these criteria, a score of 17-20 is superior, 13-16 is standard, 9-12 is under standard, and 1-8 is not acceptable.

### Production-scale honey fermentation and distillation

Two large honey fermentations were performed at Cardinal Spirits, LLC (Bloomington, IN). The first was performed on April 3, 2017 with a solution of 378 L honey and 23.2 hL H_2_O, which had an initial specific gravity of 1.0569 (14.02 Brix) and an initial pH of 4.4. Diammonium phophate (DAP, 1.05 kg; BSG Wine, Napa, CA) and Fermax (587 g; BSG HandCraft, Shakopee, MN) were added to this solution as yeast nutrients, and then a 1:1:1 mixture of YH178, YH179, and WLP715 was added to inoculate the medium. The culture was maintained at a temperature of 28.5-30°C in a stainless steel fermenter, and gravity and pH were monitored daily. After 1 day of fermentation, 1.05 kg DAP and 587 g Fermax were again added to the fermentation, and when the gravity decreased to < 6 Brix, 587 g Fermax was added one last time. The fermentation proceeded for ∼9 days, reaching a terminal gravity of 0.13 Brix (7.35% ABV) prior to distillation to approximately 150 proof.

The second large-scale fermentation was performed on September 18, 2017 with a solution of 549 L honey and 29.2 hL H_2_O, which had an initial specific gravity of 1.0708 (17.24 Brix) and an initial pH of 5.04. The same nutrient addition schedule and yeast blend as above was used. The culture was maintained at a temperature of 28.5-31°C in a stainless steel fermenter, and gravity and pH were again monitored daily. The fermentation proceeded for ∼7 days, reaching a terminal gravity of 0 Brix (9.14% ABV) prior to distillation to approximately 150 proof. In both this and the previous case, the distillate diluted to ∼80 proof and back-sweetened with 0.45 kg honey/3.78 L distillate. After resting this liquid for 1 month on fining agent to precipitate out proteins and pollen, the final product was a yellow-tinted clear liquid (visually reminiscent of white wine) and marketed as a 74.2 proof Honey Schnapps. Further details are available upon request.

### Statistical analysis

All statistical analyses were performed using GraphPad Prism 6 software. Differences between values were compared using Student’s t-test, and P-values < 0.05 were considered statistically significant.

## Acknowledgements

We thank the members of our sensory analysis panels for providing valuable flavor and aroma data, as well as members of the Bochman laboratory for critically reading this manuscript and providing feedback. This work was supported by startup funds from Indiana University, a Translational Research Pilot Grant from the Johnson Center for Entrepreneurship in Biotechnology, and funds from Wild Pitch Yeast, LLC (to MLB).

## Author Contributions

MLB, JH, and AQ conceived the study and designed the experiments. JB, MM, JH, and MLB performed the experiments, and all authors analyzed and interpreted the results. MLB wrote the manuscript. All authors read and approved the final version of the manuscript.

